# A microbial-sensory axis drives *Pseudomonas aeruginosa*-induced mechanical itch

**DOI:** 10.64898/2026.06.06.730581

**Authors:** Zhaohua Pang, Huijuan Ding, Huan Liu, Qiong Wang, Yu Wang, Qianwen Zhu, Yao Chen, Guodun Zhao, Ximin Hu, Zhenru Chen, Liqin Zhou, Ting Wang, Lefu Lan, Zhaobing Gao, Hong-Fei Zhang, Jing Feng, Fengxian Li

**Affiliations:** Department of Anesthesiology, Zhujiang Hospital of Southern Medical University, Guangzhou, 510282, China; State Key Laboratory of Chemical Biology, Shanghai Institute of Materia Medica, Shanghai, 201203, China; State Key Laboratory of Drug Research, Shanghai Institute of Materia Medica, Shanghai, 201203, China; University of Chinese Academy of Sciences, Beijing, 100000, China; School of Pharmaceutical Science and Technology, Hangzhou Institute for Advanced Study, University of Chinese Academy of Sciences, Hangzhou, 310024, China; Wuya College of Innovation, Shenyang Pharmaceutical University, Shenyang, 110016, China; Department of Burn and Plastic Surgery, Guangzhou First People’s Hospital, South China University of Technology, Guangzhou, 510000, China; Institute of Perioperative Medicine and Organ Protection, Zhujiang Hospital of Southern Medical University, Guangzhou, 510280, China; Key Laboratory of Mental Health of the Ministry of Education, Guangdong Province Key Laboratory of Psychiatric Disorders, Southern Medical University, Guangzhou, China

## Abstract

Cutaneous bacterial infections frequently elicit severe pruritus, prominently featuring alloknesis, a pathological state where innocuous touch provokes intense itch. However, the peripheral mechanisms translating microbial cues into touch-evoked pruritus remain unresolved. Here, we establish an epicutaneous *Pseudomonas aeruginosa* infection model that robustly isolates mechanical alloknesis from spontaneous scratching. We identify bacterial flagellin as the critical virulence factor driving this specific sensory modality via Toll-like receptor 5 (TLR5) activation exclusively within Calb1^+^ Aβ rapidly adapting low-threshold mechanoreceptors (RA-LTMRs). Mechanistically, pathogen-driven TLR5 signaling depletes intracellular PIP2, which suppresses KCNQ4-mediated M-currents and dismantles the biophysical brake on LTMR excitability. Our findings define a distinct microbial-neuronal axis that directly converts tactile stimuli into itch at the peripheral entry point, providing an infection-based framework for dissecting pathogen-sensory neuron interactions and uncovering precise therapeutic targets for chronic, touch-evoked pruritus.

TEASER

A bacterial flagellin-sensing touch neuron pathway converts innocuous touch into itch during skin infection.

## INTRODUCTION

Mechanical itch (alloknesis) represents a pathological sensory transformation wherein normally innocuous tactile stimuli evoke severe pruritus (*1*). While prevalent in chronic inflammatory and infectious dermatoses, the peripheral mechanisms that selectively sensitize touch pathways remain poorly defined (*2*-*4*). Recent paradigms establish that microbes can directly engage sensory neurons to drive itch. For instance,

Staphylococcus aureus drives robust spontaneous pruritus and secondary alloknesis by secreting V8 protease to activate PAR1 on TRPV1^+^ C-fiber nociceptors (*3*). However, it remains entirely unknown whether pathogens can selectively drive mechanical itch independent of ongoing spontaneous C-fiber pruriception, and if so, which peripheral mechanoreceptor subsets mediate this specific sensory modality.

*Pseudomonas aeruginosa* is a ubiquitous environmental pathogen responsible for waterborne dermatological infections, classically presenting as “hot tub folliculitis” (*5*-*7*). A hallmark clinical feature of this infection is intense pruritus that is highly exacerbated by the light friction of swimwear or clothing against the skin, a clear manifestation of alloknesis (*8*). Crucially, this touch-evoked itch often outlasts overt tissue inflammation (*9*). We reasoned that studying *P. aeruginosa*-induced cutaneous pathology could provide a pristine model to isolate the distinct mechanisms of infection-associated mechanical itch, effectively uncoupling it from the canonical inflammatory pain or spontaneous itch pathways.

The central neural circuits processing mechanical itch are increasingly well-characterized, prominently featuring a spinal relay where peripheral inputs engage Ucn3^+^ excitatory interneurons, subject to tonic suppression by NPY^+^ inhibitory networks (*10*, *11*). Yet, the peripheral initiation of this circuit during infection remains a critical knowledge gap. While recent work implicates minor nociceptive subsets (e.g., Piezo2 in MrgprA3^+^ neurons) in chemically induced alloknesis (*12*), the primary transducers of light touch are Aβ and Aδ low-threshold mechanoreceptors (LTMRs). These LTMRs classically mediate discriminative touch and mechanical allodynia (*13*, *14*). Whether and how microbial factors directly subvert the intrinsic excitability of these primary touch receptors to bypass spinal gating and drive alloknesis has not been demonstrated.

The precise tactile activation threshold of Aβ-LTMRs is biophysically constrained by the voltage-gated potassium channel subfamily q member 4 (KCNQ4). Operating at subthreshold membrane potentials, KCNQ4 generates a non-inactivating M-current that critically stabilizes the resting membrane potential and dictates the mechanical force required to evoke action potential (*15*). Indeed, genetic ablation or loss-of-function mutations in KCNQ4 eliminate this biophysical brake, directly resulting in profound tactile hypersensitivity of rapidly adapting mechanoreceptors in both mice and humans (*16*). While this establishes KCNQ4 as the master intrinsic regulator of normal touch sensation, it remains previously unexplored whether acute pathogenic cues can dynamically subvert this channel to drive infection-associated alloknesis.

Here, we identify a flagellin-TLR5 signaling axis that specifically targets Calb1^+^ Aβ rapidly adapting (RA)-LTMRs. We demonstrate that pathogen-driven TLR5 activation depletes intracellular PIP2, thereby suppressing KCNQ4 currents. This mechanism profoundly lowers the mechanosensory threshold, providing a direct molecular and cellular explanation for how a specific structural bacterial component hijacks peripheral tactile afferents to induce pathological mechanical itch.

## RESULTS

### Epicutaneous *P. aeruginosa* exposure evokes mechanical itch in mice

Exposure to contaminated water often results in *P. aeruginosa* infections, which manifest as rashes and pruritus in humans (*8*, *9*). To elucidate the mechanisms underlying this response, we employed a murine model of epicutaneous *P. aeruginosa* infection. Mice were exposed to the wild-type (WT) *P. aeruginosa* strain PAO1 solution for 1 hour (*3*), and itch-related behaviors were recorded for 5 consecutive days (Figure 1A). Interestingly, although the infection site developed rash-like dermatitis and skin thickening within 24 hours (Figure 1B-D), PAO1-exposed group did not develop significant spontaneous itch and the scratching bouts was comparable with that in the vehicle-treated controls over the observation period (Figure 1E).

**Figure 1.**
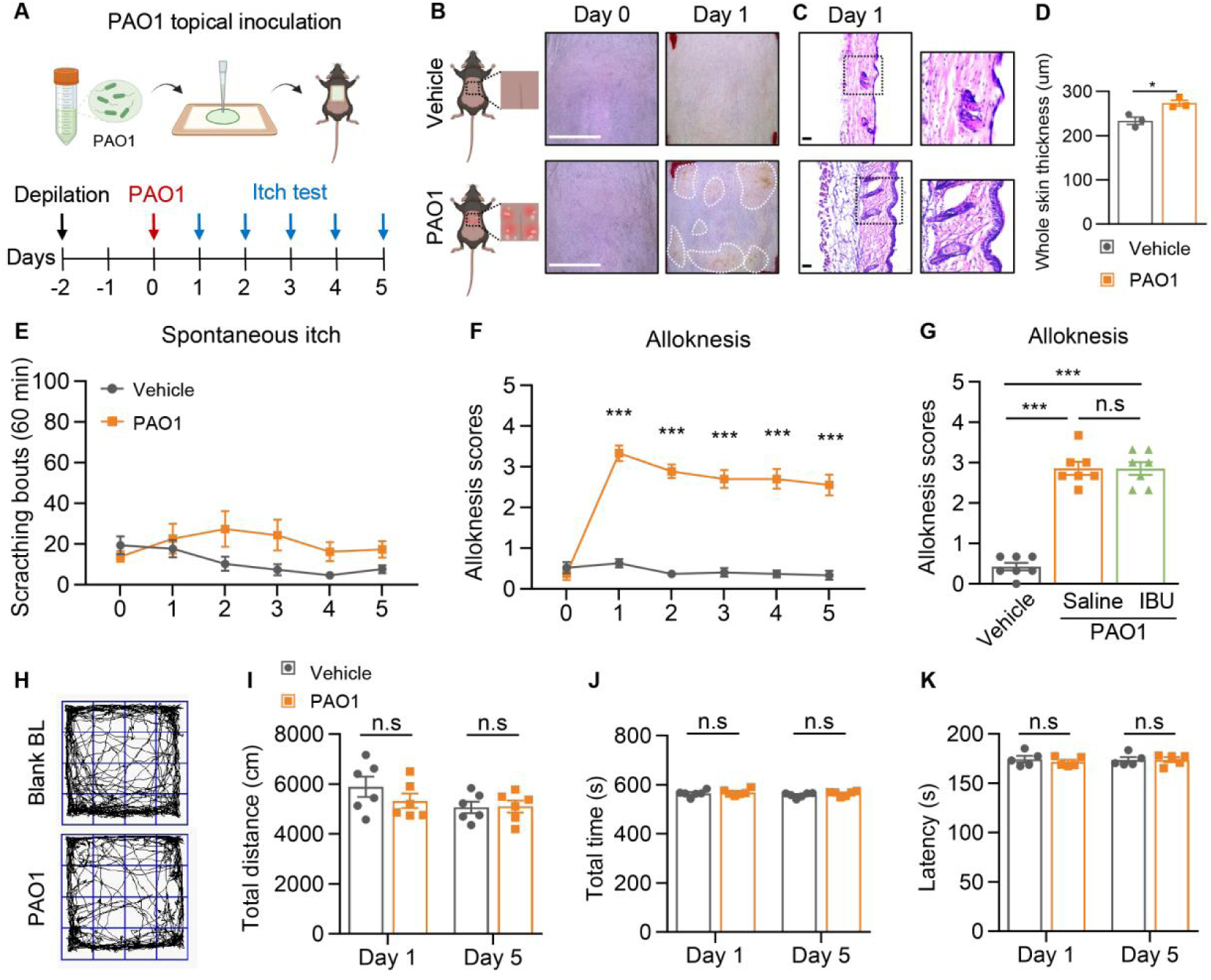
Epicutaneous *P. aeruginosa* exposure uncouples mechanical itch from spontaneous pruritus. (A) Experimental paradigm for murine epicutaneous PAO1 infection. (B) Representative macroscopic presentation of the nape skin following vehicle (LB broth) or PAO1 inoculation. Scale bar: 1 cm. (C-D) Representative H&E histopathology images (C) and quantification of epidermal thickness (D) of the nape skin following vehicle or PAO1 inoculation (n = 3 mice per group). Scale bar: 50 μm. (E-F) Spontaneous scratching bouts (E) and mechanical alloknesis scores (F) evaluated over a 5-day time course following epicutaneous exposure. (G) PAO1-induced alloknesis scores measured 1 hour post-intraperitoneal injection of saline or the NSAID ibuprofen (IBU) (n = 7 mice per group). (H) Representative open field test (OFT) movement trajectories for vehicle- and PAO1-exposed mice. (I-K) Total distance traveled (I) and ambulatory time (J) in the OFT, alongside latency to fall in the rotarod test (K) (n = 5 mice per group). Data are presented as mean ± SEM. Statistical analyses: unpaired two-tailed Student’s *t*-test (D); two-way ANOVA followed by Šídák’s multiple comparisons test (E, F, I, J, K); one-way ANOVA with Tukey’s post-hoc test (G). n.s., not significant; \**P* < 0.05, \*\*\**P* < 0.001. PAO1, WT *P. aeruginosa*; LB, Luria-Bertani.

In contrast, mechanical stimulation of the affected site with a 0.07 g von Frey filament elicited a pronounced scratching response in PAO1-treated mice as early as day 1 (Figure 1F). This response was absent in vehicle-treated controls. To confirm that the observed scratching represented itch-specific behavior rather than a pain response, we administered a nonsteroidal anti-inflammatory drug (NSAID). The alloknesis scores remained unaffected by NSAID treatment (Figure 1G), ruling out pain as the driving factor. Consistent with this interpretation, PAO1 exposure did not affect general locomotor activity or motor coordination, as assessed by open-field and rotarod tests on days 1 and 5 (Figure 1H-K). Together, these findings establish a bacteria-induced mechanical itch model in which *P. aeruginosa* selectively sensitizes the skin to innocuous touch without provoking spontaneous itch or nociceptive pain, providing a tractable platform to isolate peripheral mechanisms underlying mechanically evoked itch and alloknesis.

### Flagellin drives *P. aeruginosa*-induced mechanical itch

Flagella act as key virulence factor of *P. aeruginosa*, which participates in infection, immune activation, and biofilm formation (*17*, *18*). To elucidate the role of this component, we intradermally injected flagellin and performed behavioral assays. While higher doses (0.9 µg) of flagellin induced acute spontaneous itch and triggered mechanical pain (Figure S1), the lower dose (0.3 µg) used here failed to elicit acute spontaneous itch after intradermal injection and did not induce mechanical pain upon intraplantar delivery (Figure 2A and B), yet robustly evoked mechanically evoked itch responses (Figure 2C). These results suggest a dose-dependent and modality-specific contribution of flagellin to *P. aeruginosa*-induced mechanical itch.

**Figure 2.**
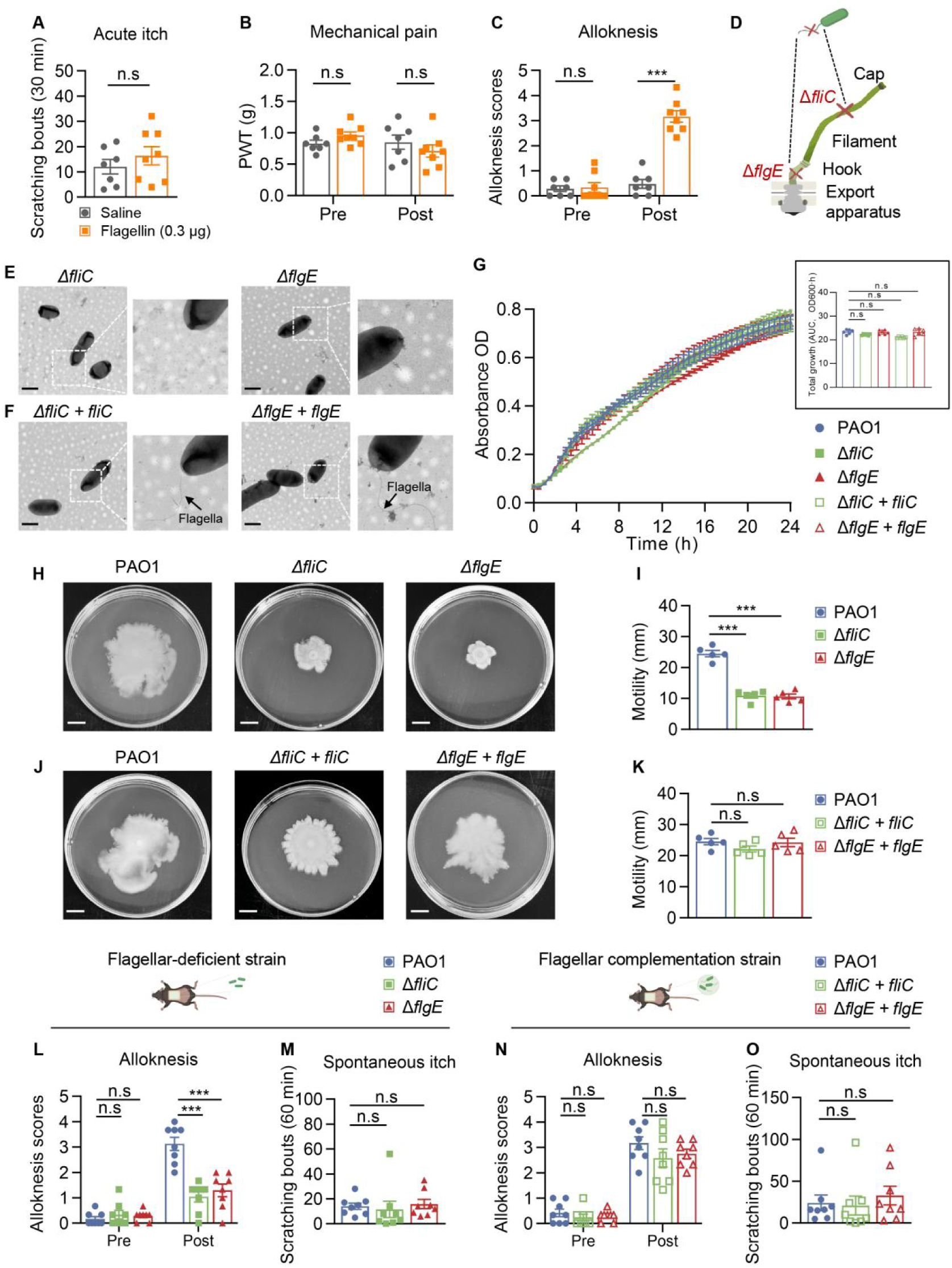
Flagellin drives *P. aeruginosa*-induced mechanical itch via flagellar structural integrity. (A-C) Acute spontaneous itch (A), mechanical pain thresholds (B), and mechanical alloknesis scores (C) following intraplantar injection of saline (n = 7 mice) or 0.3 μg purified flagellin (n = 8 mice) in WT mice. (D) Schematic detailing the genetic disruption of *P. aeruginosa* flagellar components. (E-F) Representative transmission electron micrographs of Δ*fliC* and Δ*flgE* structural mutants (E) and their respective genetically complemented strains, Δ*fliC* + *fliC* and Δ*flgE* + *flgE* (F). Scale bar: 1 μm. (G) *In vitro* growth kinetics of PAO1, mutant, and complemented strains (main panel) quantified by AUC analysis (inset). (H-K) Representative images (left) and quantification (right) of swimming motility assays on 0.3% soft agar LB plates for PAO1, structural mutants (H-I), and complemented strains (J-K) (n = 5 replicates per strain). Scale bar: 1 cm. (L-M) Mechanical alloknesis (L) and spontaneous itch (M) before and after epicutaneous inoculation with PAO1, Δ*fliC*, or Δ*flgE* strains (n = 8 mice per group). (O-P) Mechanical alloknesis (O) and spontaneous itch (P) before and after epicutaneous inoculation with PAO1 or the complemented strains (n = 8 mice per group). Data are presented as mean ± SEM. Statistical analyses: unpaired two-tailed Student’s *t*-test (A); one-way ANOVA with Tukey’s post-hoc test (G, I, K, M, O); two-way ANOVA followed by Šídák’s multiple comparisons test (B, C); two-way ANOVA with Tukey’s post-hoc test (L, N). n.s., not significant; \*\*\**P* < 0.001. See also Figure S1.

To directly assess the requirement of an intact bacterial flagellum in mechanical itch, we adopted a complementary loss- and rescue-genetic strategy targeting distinct structural components of the *P. aeruginosa* flagellum. Specifically, we generated a Δ*fliC* mutant lacking the flagellin filament subunit itself, and a Δ*flgE* mutant lacking the flagellar hook protein essential for flagellum assembly (Figure 2D), both of which disrupt flagellar structure without broadly affecting bacterial viability. Transmission electron microscopy confirmed the absence of intact flagella in both Δ*fliC* and Δ*flgE* strains (Figure 2E), accompanied by a marked loss of flagellum-dependent swimming motility (Figure 2H and I), while growth kinetics remained comparable to the wild type *P. aeruginosa* strain PAO1 (Figure 2G), indicating preserved viability and proliferation. To ensure specificity, we reintroduced *fliC* or *flgE* into the corresponding mutant strains, which restored flagellar architecture and motility (Figure 2F, J, K).

Functionally, epicutaneous exposure to either Δ*fliC* or Δ*flgE* mutants resulted in a profound reduction in mechanically evoked itch compared with PAO1 (Figure 2L and M). Strikingly, genetic complementation of *fliC* or *flgE* fully rescued the ability of these strains to induce mechanical alloknesis *in vivo* (Figure 2N and O). Together, these loss- and rescue-experiments establish that an intact bacterial flagellum is required for *P. aeruginosa*-induced mechanical itch, supporting a specific role for flagellum-dependent signaling rather than nonspecific effects of bacterial burden or growth. Collectively, these findings establish that flagellin is both necessary and sufficient to mediate *P. aeruginosa*-induced mechanical itch.

### Flagellin-induced mechanical itch is mediated by neuronal TLR5 signaling

While the major bacterial component lipopolysaccharide (LPS) is known to influence host immune and sensory responses (*19*, *20*), global deletion of its canonical receptor toll-like receptor 4 (TLR4) did not alter flagellin/PAO1-evoked alloknesis compared with control mice (Figure S2). In contrast, Toll-like receptor 5 (TLR5) serves as the canonical receptor for bacterial flagellin and activates MyD88-dependent signaling cascades upon engagement (*21*-*23*). To test whether TLR5 is required for flagellin-evoked mechanical itch *in vivo*, we intradermally injected flagellin into the nape and observed a marked reduction in alloknesis in *Tlr5-*deficient mice relative to littermate controls (Figure 3A and B). Consistently, *P. aeruginosa*-induced mechanical itch was also significantly attenuated in *Tlr5*-deficient mice (Figure 3C and D), establishing TLR5 as a critical host sensor linking bacterial exposure to mechanically evoked itch.

**Figure 3.**
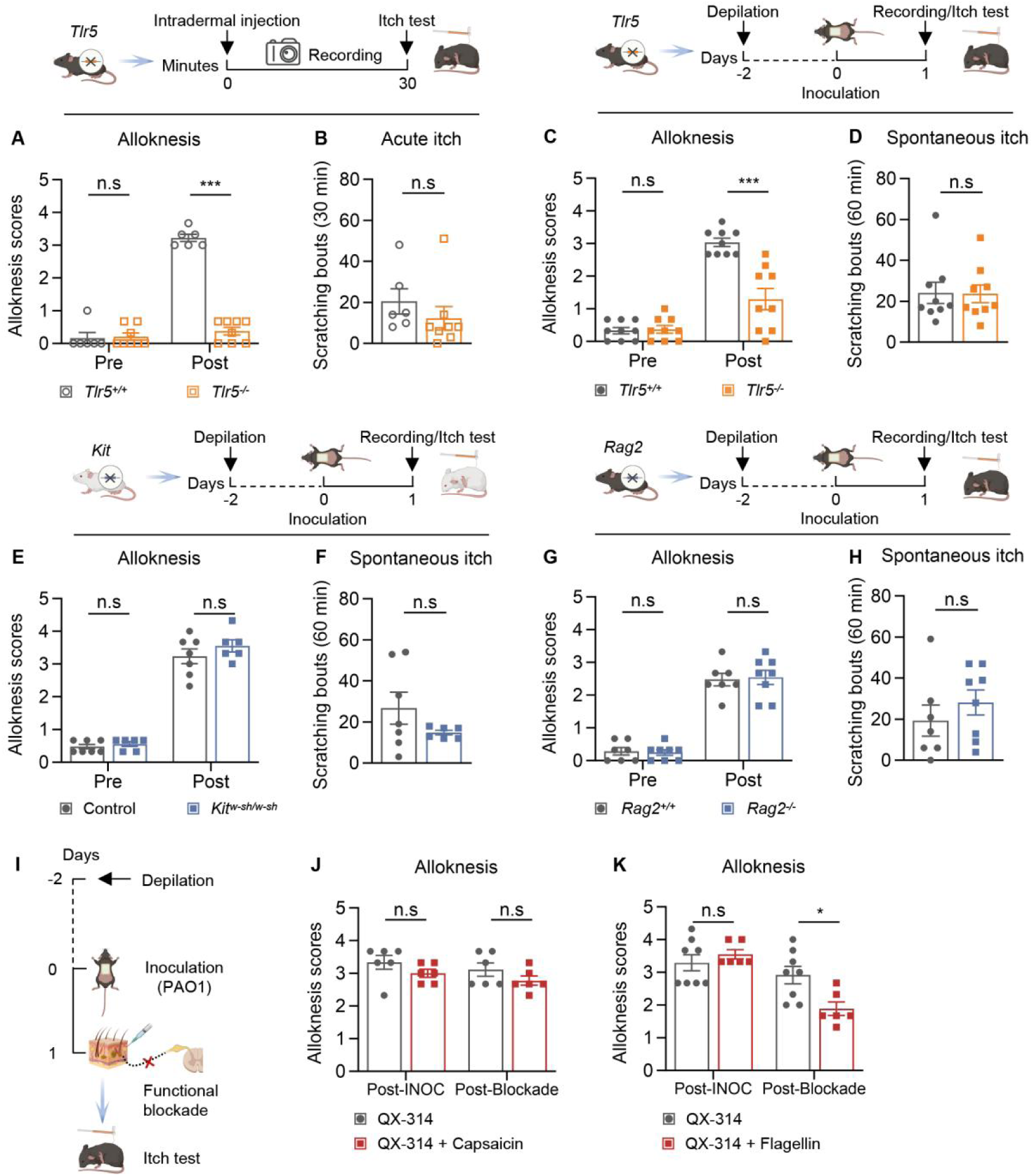
Flagellin-evoked alloknesis requires neuronal, but not immune, TLR5 signaling. (A-B) Mechanical alloknesis (A) and acute spontaneous itch (B) following intradermal injection of 0.3 μg flagellin in *Tlr5^-/-^* (n = 8 mice) and *Tlr5^+/+^* littermates (n = 6 mice). (C-D) Mechanical alloknesis (C) and spontaneous itch (D) before and after PAO1 epicutaneous exposure in *Tlr5^-/-^* and *Tlr5^+/+^* littermates (n = 9 mice per genotype). (E-F) Mechanical alloknesis (E) and spontaneous itch (F) in mast cell-deficient *Kit^W-sh/W-sh^* (n = 6 mice) and WT littermates (n = 7 mice) exposed to PAO1. (G-H) Mechanical alloknesis (G) and spontaneous itch (H) in T/B cell-deficient *Rag2^-/-^* (n = 8 mice) and *Rag2^+/+^* littermates (n = 7 mice) exposed to PAO1. (I) Schematic of the targeted pharmacological silencing paradigm using QX-314 entry via modality-specific ion channels. (J-K) PAO1-induced alloknesis following intraplantar co-injection of QX-314 with the TRPV1 agonist capsaicin (J, n = 6 mice per group) or the TLR5 agonist flagellin (K, n = 8 and 6 mice, respectively). Data are presented as mean ± SEM. Statistical analyses: two-way ANOVA followed by Šídák’s multiple comparisons test (A, C, E, G, J, K); unpaired two-tailed Student’s *t*-test (B, D, F, H). n.s., not significant; \**P* < 0.05, \*\*\**P* < 0.001. INOC, inoculation. See also Figure S2.

Because TLR5 is expressed across multiple cell types, including epithelial and immune compartments (*24*, *25*), we next asked whether the behavioral phenotype reflects indirect immune-to-neuron signaling or a neuron-intrinsic pathway. Using genetic and pharmacologic dissection, we found that mast cell deficiency or T/B-cell deficiency did not impact PAO1-evoked alloknesis (Figure 3E-H). In contrast, while acute silencing of TRPV1-expressing C-type high-threshold mechanoreceptors (HTMRs) with capsaicin/QX-314 (*26*) did not alter mechanically evoked itch responses (Figure 3I and J), silencing of TLR5-expressing A-type LTMRs with flagellin/QX-314 (*13*) robustly suppressed alloknesis following PAO1 exposure (Figure 3K). Together, these results support a model in which bacterial flagellin drives mechanical itch predominantly through TLR5-dependent signaling in peripheral sensory neurons rather than via canonical immune cell-dependent itch pathways.

### TLR5 is abundantly expressed on Calb1-positive and Ntrk2-positive LTMRs

To elucidate the cellular basis of TLR5-dependent mechanical itch in the *P. aeruginosa* model, we re-analyzed single-cell RNA-seq datasets of dorsal root ganglion (DRG) neurons (Figure S3) (*27*). This analysis revealed that *Tlr5* is abundantly expressed in two distinct subsets of LTMRs: *Ntrk2*-expressing Aδ-LTMRs, which are medium-diameter, thinly myelinated neurons that innervate hair follicles with circumferential endings and respond to light dynamic touch (*14*), and *Calb1*-expressing Aβ RA-LTMRs, which are large-diameter, heavily myelinated neurons responsible for fast-conducting tactile signals via Merkel and Meissner corpuscles (*28*, *29*).

To functionally label these populations, we used tamoxifen-inducible *Ntrk2-CreER* and trimethoprim (TMP)-inducible *Calb1-dgCre* mice crossed with *Ai9* reporter mice, allowing visualization of respective neuronal subtypes (Figure S4). In agreement with transcriptomic data, *Tlr5* expression showed minimal overlap with nociceptive TRPV1^+^ small-diameter neurons or TH^+^ C-LTMRs (Figure 4A-D), but predominantly colocalized with NF200^+^ myelinated neurons (Figure 4E and F). Quantification revealed that 40.18% of TLR5^+^ neurons were Calb1-positive (Figure 4G and H) and 48.95% were Ntrk2-positive (Figure 4I and J), highlighting a potential role for *Tlr5*-expressing LTMRs in mediating mechanotransduction in bacterial-induced itch.

**Figure 4.**
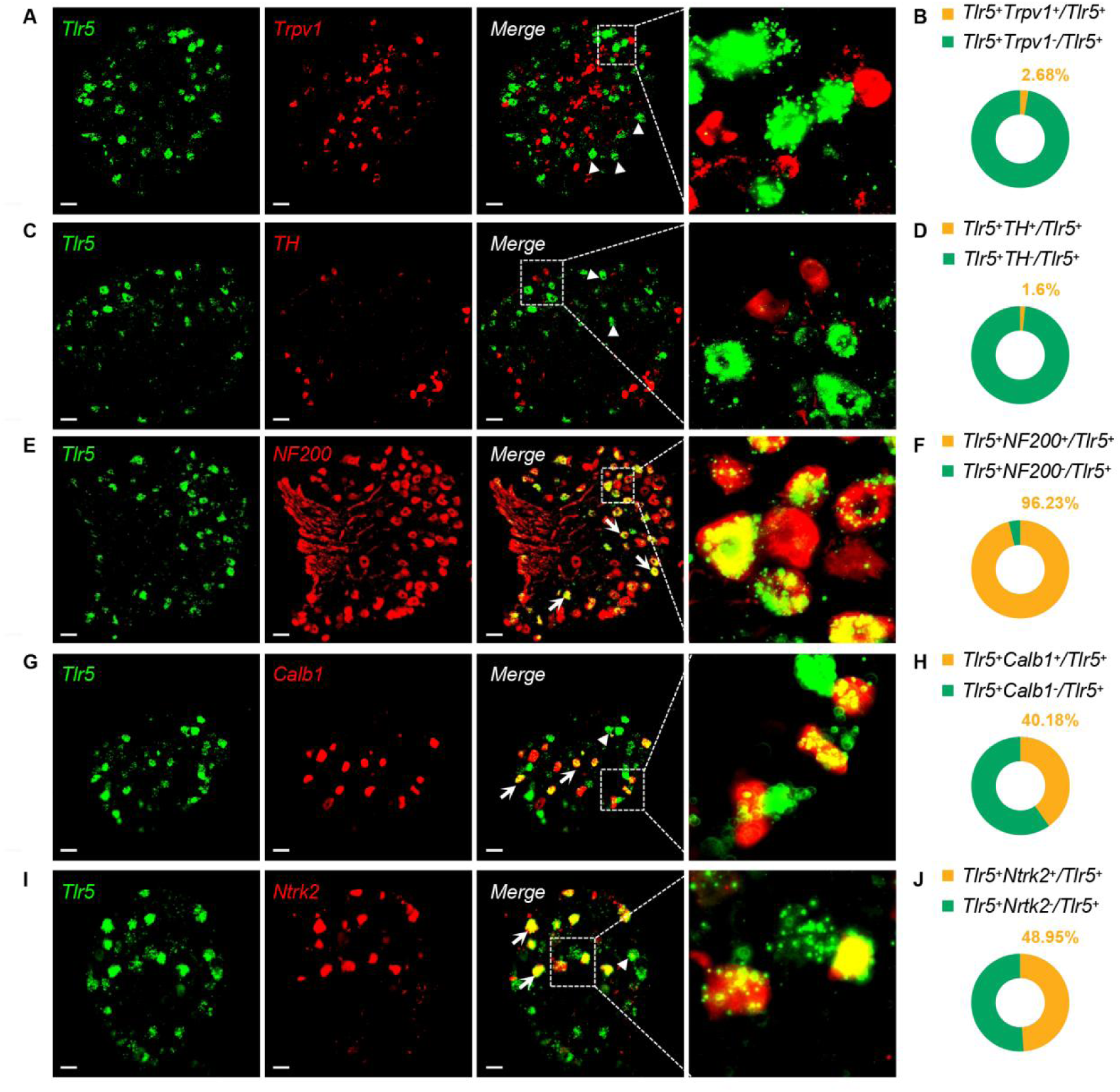
TLR5 is highly enriched in distinct Calb1^+^ and Ntrk2^+^ LTMR subpopulations. (A, C, E, G, I) Multiplexed *in situ* hybridization (ISH) for *Tlr5* mRNA (green) coupled with immunofluorescence (IF) for distinct neuronal subset markers (red) in whole-mount mouse dorsal root ganglia (DRG). Evaluated markers include Trpv1 (A), Tyrosine Hydroxylase (TH) (C), NF200 (E), Calb1 (G), and Ntrk2 (I). White arrows denote cells co-expressing *Tlr5* and the specified marker; white arrowheads denote neurons exclusively expressing *Tlr5*. Scale bar: 50 μm. (B, D, F, H, J) Quantitative analysis of the percentage of TLR5^+^ neurons co-expressing the indicated neuronal markers. See also Figure S3-4.

### Calb1-positive Aβ LTMRs as peripheral initiators of flagellin-induced mechanical itch

Large-scale transcriptomic and functional profiling studies have established that LTMRs are molecularly and physiologically heterogeneous, comprising distinct Aβ and Aδ subtypes with specialized roles in touch encoding and mechanical hypersensitivity (*10*, *14*, *30*). In particular, *Calb1*-expressing Aβ RA-LTMRs have been implicated in fine tactile discrimination and dynamic touch processing (*29*), whereas *TrkB*/*Ntrk2*-expressing Aδ-LTMRs preferentially contribute to mechanical pain hypersensitivity (*14*). Despite these advances, whether defined LTMR subsets directly participate in mechanical itch, especially in pathological contexts such as infection, has remained unclear.

To address this gap, we first established genetic tools to selectively manipulate defined LTMR subsets in adult mice. Calb1-positive Aβ RA-LTMRs were targeted using an intersectional *Calb1-dgCre* driver line, in which Cre recombinase expression is inducible by TMP, allowing temporal control of recombination in mature sensory neurons. *Calb1-dgCre* mice were crossed with Cre-dependent hM3Dq-DREADD reporter mice to allow chemogenetic activation of this population (Figure 5A). Activation of Calb1^+^ neurons induced minimal spontaneous scratching but robustly potentiated mechanically evoked itch without eliciting nocifensive behaviors (Figure 5B-D), indicating that activation of this LTMR subset is sufficient to bias tactile input toward itch rather than pain. In contrast, chemogenetic activation of Ntrk2^+^ Aδ-LTMRs, achieved by crossing *Ntrk2-CreER* with the same reporter line (Figure 5E), selectively enhanced mechanical pain responses without inducing mechanical or spontaneous itch (Figure 5F-H), consistent with their established role in mechanical hyperalgesia.

**Figure 5.**
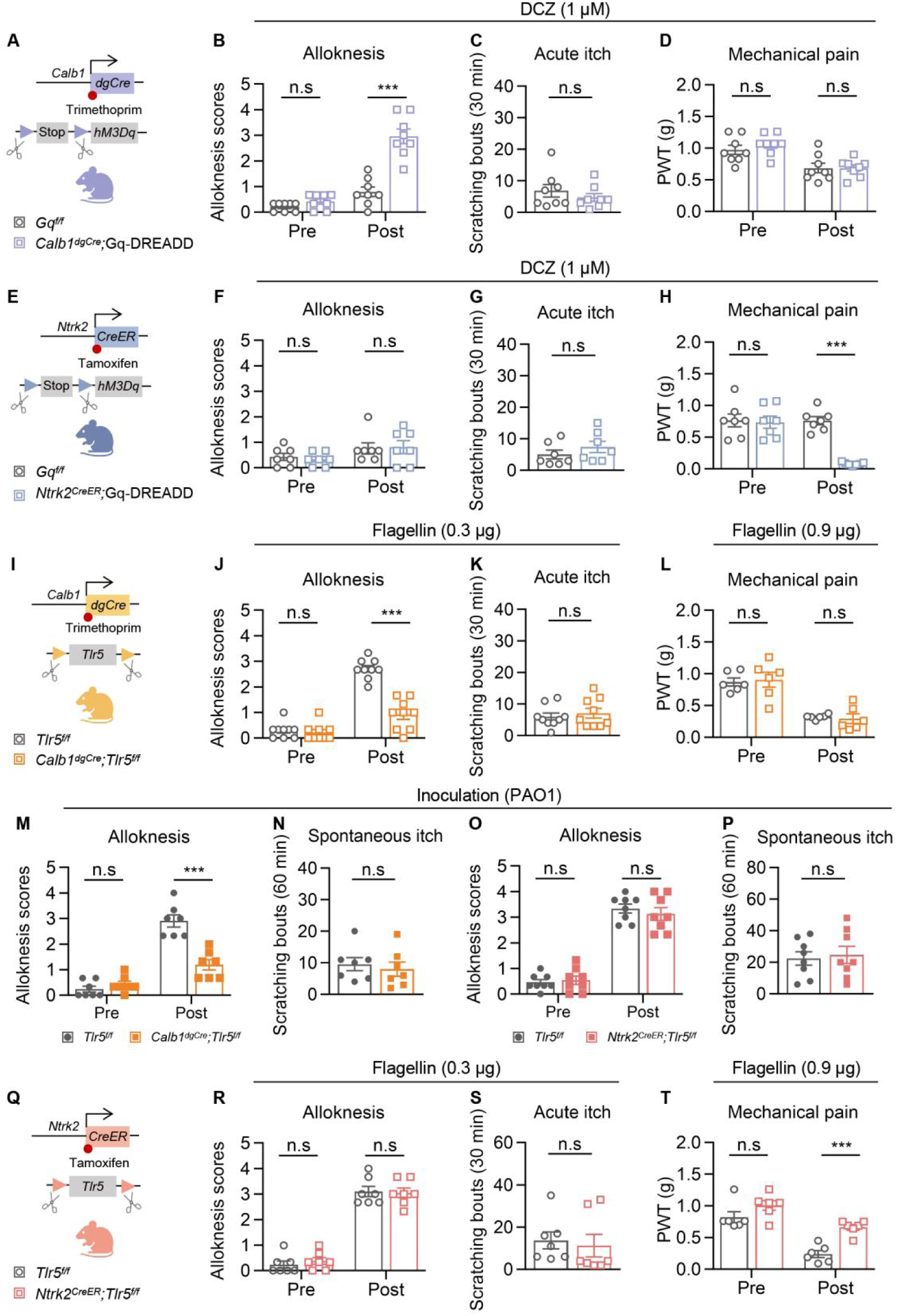
Calb1^+^ Aβ RA-LTMRs selectively govern flagellin-induced mechanical itch. (A) Intersectional genetic strategy for selective Gq-DREADD-mediated chemogenetic activation of Calb1^+^ neurons. (B-D) Behavioral parameters including alloknesis (B), acute spontaneous itch (C), and mechanical pain thresholds (D) evaluated before and after intraplantar DCZ (1 μM) application in *Calb1^dgCre^;*Gq-DREADD (n = 8 mice) versus control mice (n = 8 mice). (E) Intersectional genetic strategy for selective chemogenetic activation of Ntrk2^+^ neurons. (F-H) Behavioral parameters including alloknesis (F), acute spontaneous itch (G), and mechanical pain thresholds (H) evaluated before and after intraplantar DCZ (1 μM) application in *Ntrk2^CreER^;*Gq-DREADD (n = 7 mice) versus control mice (n = 7 mice). (I) Strategy for conditional ablation of TLR5 in Calb1^+^ neurons (*Calb1^dgCre^;Tlr5^f/f^*). (J-L) Assessment of alloknesis (J), acute spontaneous itch (K) following 0.3 μg flagellin injection, and mechanical pain thresholds (L) following 0.9 μg flagellin injection in *Calb1^dgCre^;Tlr5^f/f^* mice versus *Tlr5^f/f^* littermates (n = 6-9 mice per group). (M-N) Mechanical alloknesis (M) and spontaneous itch (N) before and after PAO1 exposure in *Calb1^dgCre^;Tlr5^f/f^* mice and controls (n = 7 mice per group). (O-P) Mechanical alloknesis (O) and spontaneous itch (P) before and after PAO1 exposure in *Ntrk2^CreER^;Tlr5^f/f^* mice and controls (n = 8 mice per group). (Q) Strategy for conditional ablation of TLR5 in Ntrk2^+^ neurons (*Ntrk2^CreER^;Tlr5^f/f^*). (R-T) Assessment of alloknesis (R), acute spontaneous itch (S) following 0.3 μg flagellin injection, and mechanical pain thresholds (T) following 0.9 μg flagellin injection in *Ntrk2^CreER^;Tlr5^f/f^* mice versus *Tlr5^f/f^* littermates (n = 6-7 mice per group). Data are presented as mean ± SEM. Statistical analyses: two-way ANOVA followed by Šídák’s multiple comparisons test (B, D, F, H, J, L, M, O, R, T); unpaired two-tailed Student’s *t*-test (C, G, K, N, P, S). n.s., not significant; \*\**P* < 0.01, \*\*\**P* < 0.001. DCZ, deschloroclozapine.

We next interrogated the role of TLR5 signaling within these LTMR subsets. Selective deletion of TLR5 in Calb1^+^ neurons using the *Calb1^dgCre^;Tlr5^f/f^*mice markedly attenuated both flagellin-evoked and POA1-induced mechanical itch, while leaving spontaneous itch and mechanical pain intact (Figure 5I-N). By contrast, deletion of TLR5 in Ntrk2^+^ neurons failed to affect mechanical alloknesis, despite reducing high-dose flagellin-induced mechanical pain (Figure 5O-T). Together, these findings identify Calb1^+^ Aβ RA-LTMRs as the principal peripheral sensory neurons that detect bacterial cues and initiate mechanical itch.

Together, these data reveal a role for Calb1^+^ Aβ-LTMRs in infection-associated mechanical itch. Unlike nociceptive C-fibers that mediate spontaneous itch or Aδ-LTMRs that contribute to mechanical pain, Calb1^+^ Aβ-LTMRs serve as a dedicated peripheral entry point through which microbial signaling selectively transforms innocuous touch into itch.

### Intracellular TLR5-KCNQ4 signaling modulates excitability of LTMR neurons

KCNQ4 expression is enriched across multiple LTMR populations, including Aδ-LTMRs, Aβ RA-LTMRs, and Aβ field-LTMRs (*31*), consistent with prior observations that KCNQ4 protein localizes to lanceolate and circumferential endings in the skin. Given that KCNQ4 is a principal subunit underlying M-type potassium currents that restrain excitability in mechanosensory neurons (*16*, *32*), we hypothesized that TLR5 activation may modulate LTMR excitability through KCNQ4. Consistent with this idea, reanalysis of single-cell RNA-seq datasets from DRG neurons (*27*) revealed strong co-expression of *Kcnq4* and *Tlr5* within LTMR subsets, with prominent enrichment in Aδ-LTMRs and Aβ RA-LTMRs (Figure S3).

We further validated this molecular overlap using RNA in situ hybridization and immunofluorescence, which showed that 64.99% of TLR5^+^ neurons expressed *Kcnq4* (Figure 6A and B). Consistently, FACS-sorted TLR5^+^ neurons from *Tlr5^Cre^;Ai9* mice exhibited the highest *Kcnq4* expression among *Kcnq*2-5 family members by RT-qPCR (Figure 6C). Retrograde tracing with Cholera Toxin Subunit B (CTB) injected into the nape skin of *Tlr5^Cre^;Ai9* mice further revealed that 88.24% of cutaneous-projecting TLR5^+^ neurons co-expressed *Kcnq4* (Figure 6D and E), highlighting preferential enrichment of *Kcnq4* within mechanosensory TLR5^+^ afferents.

**Figure 6.**
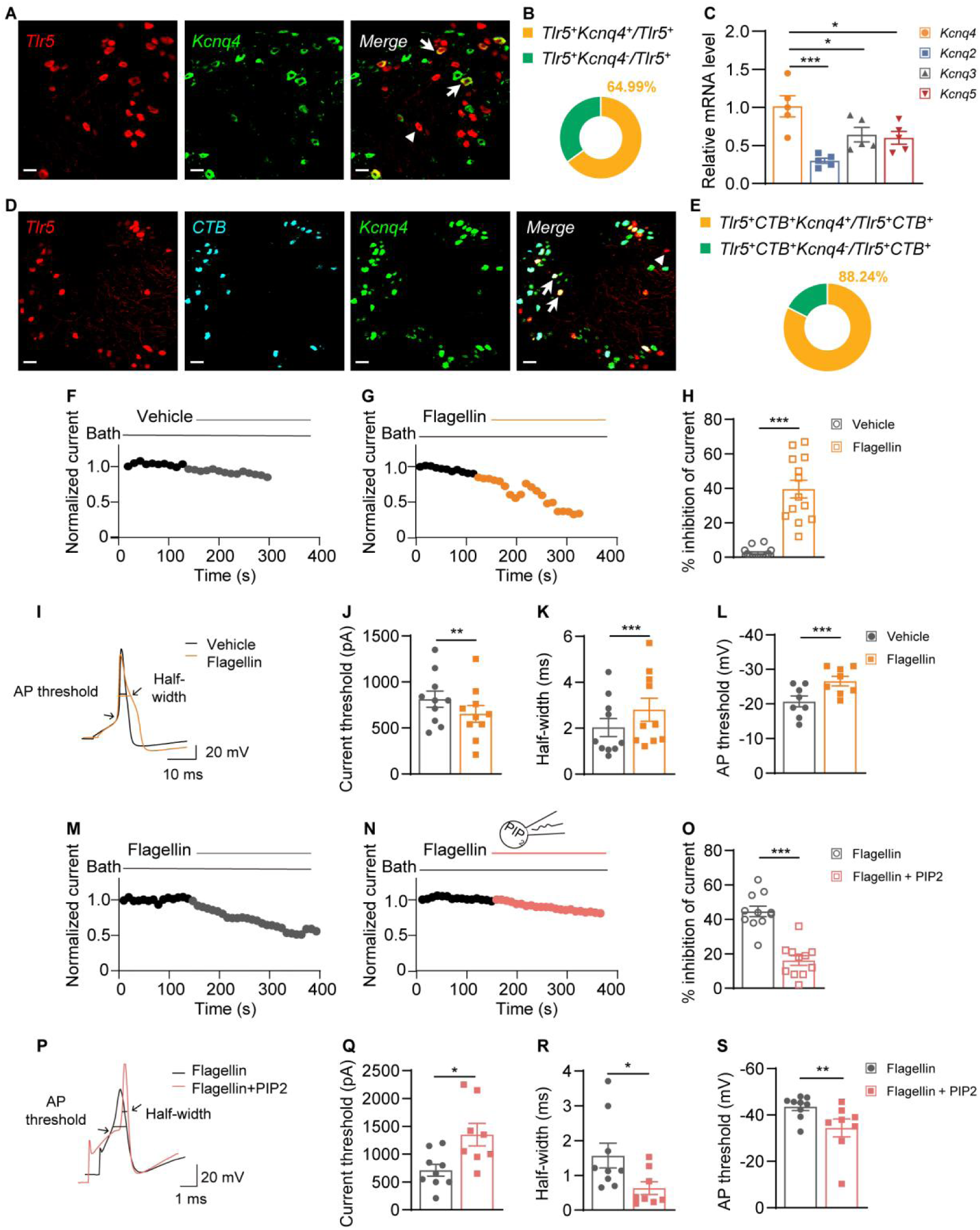
Flagellin-TLR5 signaling actively enhances LTMR excitability via PIP2-dependent KCNQ4 suppression. (A) Representative co-expression analysis utilizing IF (*Tlr5-tdTomato*, red) and mRNA ISH (*Kcnq4*, green) in naïve mouse DRG. Scale bar: 50 μm. (B) Quantification of the percentage of Tlr5^+^ neurons co-expressing *Kcnq4*. (C) RT-qPCR validation of *Kcnq* family channel transcript enrichment in FACS-sorted TLR5^+^ neurons. (D) Retrograde tracing analysis mapping cutaneous projections; co-localization of TLR5 (red), peripherally injected Alexa Fluor 647-CTB (cyan), and *Kcnq4* (green) in DRG neurons. (E) Quantification of the percentage of skin-projecting (CTB^+^) TLR5^+^ neurons expressing *Kcnq4*. (F-H) Representative voltage-clamp recordings of M-currents (F, G) and quantified inhibition percentages (H) in acutely dissociated TLR5^+^ DRG neurons exposed to vehicle or 50 nM flagellin. (I-L) Representative current-clamp recordings displaying action potential firing patterns in response to step current injections (I), with quantitative analyses of rheobase (J), action potential half-width (K), and voltage threshold for action potential firing (L) under vehicle or flagellin treatment. (M-O) Representative M-current traces (M, N) and inhibition quantification (O) demonstrating that intracellular dialysis with PIP2 (200 μM) rescues flagellin-induced M-current suppression. (P-S) Representative current-clamp recordings (P) and quantitative electrophysiological properties including rheobase (Q), half-width (R), and voltage threshold (S) demonstrating that PIP2 supplementation normalizes intrinsic neuronal excitability despite flagellin exposure. Data are presented as mean ± SEM. Statistical analyses: one-way ANOVA with Tukey’s post-hoc test (C); unpaired two-tailed Student’s *t*-test (H-L, O-S). n.s., not significant; \**P* < 0.05, \*\*\**P* < 0.001. See also Figure S5.

To test whether TLR5 activation functionally suppresses KCNQ4-mediated M-currents, we performed whole-cell patch-clamp recordings. In acutely dissociated TLR5^+^ DRG neurons, application of 50 nM flagellin reduced M-current amplitude by 40.03% (Figure 6F-H). Similarly, in Chinese Hamster ovary (CHO) cells co-expressing *Tlr5* and *Kcnq4*, flagellin inhibited M-currents by 57.75% (Figure S5A-C). Consistent with M-current suppression, flagellin-treated TLR5^+^ DRG neurons exhibited reduced rheobase, broadened action potential half-width, and lowered firing thresholds (Figure 6I-L), indicating increased intrinsic excitability.

Because KCNQ4 channel activity critically depends on membrane phosphatidylinositol 4,5-bisphosphate (PIP2) (*33*), and TLR5 signaling has been reported to engage PI3K-PLC pathways capable of depleting PIP2 (*34*), we hypothesized that PIP2 loss mediates TLR5-dependent M-current suppression. Indeed, intracellular supplementation with 200 µM PIP2 fully rescued flagellin-induced M-current inhibition in both TLR5^+^ DRG neurons (Figure 6M-O) and CHO cells co-expressing *Tlr5* and *Kcnq4* (Figure S5D-F). PIP2 supplementation also normalized electrophysiological parameters altered by flagellin, including rheobase, action potential width, and firing threshold (Figure 6P-S).

Together, these results support a mechanistic model in which bacterial flagellin activates neuronal TLR5 to trigger PIP2 depletion and suppress KCNQ4-dependent M-currents, thereby enhancing the excitability of Calb1^+^ Aβ-LTMRs. This pathogen-driven modulation of a core excitability regulator provides a direct molecular link between microbial sensing in the skin and the emergence of mechanical itch during *P. aeruginosa* infection.

## DISCUSSION

The fundamental conceptual advance of this study lies in uncoupling mechanical itch from canonical nociceptive and chemical pruriceptive pathways during bacterial infection. While current paradigms emphasize that microbes drive spontaneous itch through the activation of polymodal C-fibers, exemplified by the Staphylococcus aureus V8 protease-PAR1 axis in TRPV1^+^ neurons (*3*), our findings establish a distinct and parallel infectious mechanosensory axis. We demonstrate that *Pseudomonas aeruginosa* selectively hijacks Calb1^+^ RA-LTMRs via structural flagellin to drive alloknesis. This defines a framework wherein disparate microbial virulence factors differentially partition somatosensory circuits: secreted proteases engage unmyelinated pruriceptors to evoke spontaneous scratching, whereas structural appendages engage myelinated mechanoreceptors to convert innocuous touch into pathological itch.

Identifying Calb1^+^ Aβ RA-LTMRs as the dedicated peripheral initiators for microbe-induced mechanical itch resolves a critical upstream gap in pruriceptive neurobiology. Extensive circuit-mapping has delineated a spinal architecture where Ucn3^+^ excitatory interneurons act as the central relay for mechanically evoked itch, gated by NPY^+^ inhibitory networks (*10*, *11*). However, the specific peripheral afferents that convey disease-relevant signals to access these spinal pathways during pathology remained obscured. By demonstrating that TLR5^+^/Calb1^+^ Aβ RA-LTMRs are both necessary and sufficient for infection-evoked alloknesis, we identify the specific cutaneous afferents that capture microbial signals, providing the requisite upstream sensory drive that ultimately interfaces with these established central itch networks.

Mechanistically, our data reveal that this host-pathogen interaction converges on the biophysical modulation of the KCNQ4 channel. In physiological states, KCNQ4-mediated M-currents function as an intrinsic brake, dictating the high tactile thresholds of rapidly adapting mechanoreceptors (*16*). We demonstrate that pathogen-driven TLR5 activation acts as a peripheral excitability switch, depleting membrane PIP2 to extinguish KCNQ4 currents. This actively subverts the biophysical constraints of the mechanosensory neuron, lowering its activation threshold and rendering the afferent hypersensitive. This pathogen-driven suppression of KCNQ4 represents a pathological extension of its physiological role, adding definitive molecular depth to the skin-to-spinal sensory pathway.

Despite the interesting findings, the study is not without limitations. First, although our data identify Calb1^+^ Aβ RA-LTMRs as the peripheral sensory neurons that initiate infection-associated mechanical itch, the downstream spinal circuits through which these neurons engage itch-processing networks remain to be determined. Second, our findings were obtained primarily in mouse models of *P. aeruginosa* infection, and whether a similar TLR5-KCNQ4 signaling mechanism operates in human sensory neurons will require further investigation. Future studies addressing these questions will help define how microbial signals are integrated into somatosensory circuits across species and disease contexts.

By defining this flagellin-TLR5-KCNQ4 signaling axis, this study elucidates how structural microbial components selectively alter the intrinsic excitability of mechanosensory neurons (Figure S6). Targeting this specific peripheral lipid-ion channel microenvironment offers a precise therapeutic strategy for chronic, touch-evoked pruritus, sidestepping the broader sensory deficits associated with global tactile or nociceptive inhibition.

## MATERIALS AND METHODS

### Experimental model and subject details Mouse models

#### Animal maintenance

All animal experiments were conducted in accordance with the protocols approved by the Institutional Animal Care and Use Committee (IACUC) of the Shanghai Institute of Materia Medica, Chinese Academy of Sciences, and adhered to the ethical guidelines of the International Association for the Study of Pain. Both adult male and female mice (8-12 weeks old; 20-25 g body weight) were used, and no sex differences in itch and pain responses were observed across experimental groups. C57BL/6 wild-type mice were purchased from Shanghai SLAC laboratory Animal Co, Ltd (Shanghai, China). *Tlr5*^-/-^ (Stock No: NM-KO-18059 from Shanghai Model Organisms Center), *Tlr4*^-/-^ (Stock No: C001234 from Cyagen) and *Rag2^-/-^*(Stock No: C001324 from Cyagen). *Kit^W-sh/W-sh^* mice were kindly provided by by the Liu Lab. To identify Tlr5-positive, Calb1-positive, and TrkB-positive neurons, *Tlr5^Cre^* mice (Stock No: NK-KI-215034 from Shanghai Model Organisms Center), *Calb1^dgCre^* (*Calb1^DHFR-EGFP-Cre^*, Stock No: 8235517 from Cyagen) and *Ntrk2^CreER^* (Stock No: NM-KI-200146 from Shanghai Model Organisms Center) mice were crossed with the reporter mice *ROSA26^Tdtomato^* (*Ai9*) (Stock No: T002249 from Gempharmatech), respectively. *Calb1^dgCre^;*Gq-DREADD, and *Ntrk2^CreER^;*Gq-DREADD were obtained by crossing *Calb1^dgCre^*or *Ntrk2^CreER^* mice with *R26*-LSL-Gq-DREADD (Stock No: 026220 from Jackson Laboratory), respectively. Conditional deletion of *Tlr5* in Calb1- or TrkB-positive neurons was achieved by crossing *Tlr5^f/f^* (Stock No: NM-CKO-231637 from Shanghai Model Organisms Center) with a *Calb1^dgCre^*, *Ntrk2^CreER^*mouse line, respectively. To induce Cre activity, *Calb1^dgCre^* mice were treated with intraperitoneal injection of freshly prepared Trimethoprim (TMP; 150 mg/kg body weight; Sigma-Aldrich, Cat# T78883) for three consecutive days (*31*, *35*). TMP was dissolved sequentially by adding DMSO, PEG300, Tween-80, and saline. *Ntrk2^CreER^* mice were treated with intraperitoneal injection of freshly prepared tamoxifen (100 mg/kg body weigh; Macklin, Cat# C16406746) dissolved in corn oil for five consecutive days. Mice were housed under standard 12/12-h light/dark cycle conditions at a constant temperature of 23 ± 1℃ with free access to food and water throughout the experiments.

### *P. aeruginosa* skin infection model

*P. aeruginosa*-induced alloknesis murine model was established as previously described (*3*). Three days prior to infection, the dorsal skin of each mouse was depilated using a mild depilatory cream. Mice were then habituated to a red acrylic box (10 cm × 10 cm × 10 cm) for 4 h daily over three consecutive days. On the day of infection, mice were anesthetized with sevoflurane. A bacterial suspension (120 μL; OD600 = 5-6, equivalent to 5-6 × 10⁸ CFU/mL) was affixed to the cotton pad of a transparent dressing (Tegaderm™; 3M), which was then attached to the depilated dorsal skin. Mice were subsequently returned to the boxes, and the dressing was removed under anesthesia after 1 h of exposure. Mice that dislodged the dressing during this period were excluded. Unless otherwise indicated, alloknesis and spontaneous itch behavioral tests were performed on day 1 post-inoculation.

### Bacteria and cell line details

#### *Pseudomonas aeruginosa* PAO1 strains information

The bacterial strains and plasmids used in this study are listed in Table S1. Unless otherwise noted, *P. aeruginosa* PAO1 (*36*) and its derivatives were grown in Luria-Bertani (LB) broth (Becton Dickinson, San Jose, CA, USA). *Escherichia coli* (*E. coli*) were cultured in LB broth. All cultures were incubated at 37℃ with shaking at 250 revolutions per minute (rpm). For plasmid maintenance, antibiotics (Sangon Biotech, Shanghai, China) were used at the following concentrations: for *P. aeruginosa*, carbenicillin at 150 µg/mL in LB medium; for *E. coli*, carbenicillin at 100 µg/mL and gentamicin at 10 µg/mL in LB medium.

#### Construction of Δ*fliC* and Δ*flgE* mutants

To generate the *fliC* null mutant (Δ*fliC*), polymerase chain reactions (PCRs) were performed to amplify sequences upstream and downstream of *fliC*. The upstream fragment was amplified from PAO1 genomic DNA using primers D-*fliC*-up-F and D-*fliC*-up-R, while the downstream fragment was amplified with primers D-*fliC*-down-F and D-*fliC*-down-R. The vector pEX18Ap (*37*) was assembled with the upstream and the downstream regions of the *fliC* gene using MultiF Seamless Assembly Mix (ABclonal Biotechnology, Wuhan, China), yielding pEX18Ap::*fliCUD*. A 1.8 kb gentamicin resistance cassette cut from pPS858 with BamHI was cloned into pEX18Ap::*fliCUD*, yielding pEX18Ap::*fliCUGD* (*37*). The assembled vector was transformed to *E. coli* DH5α for amplification by heat shock at 42℃ and sequenced to ensure that no unwanted mutations resulted. The plasmid for gene replacement was conjugally transferred from *E.coli* S17 λ-pir into PAO1 and selected for gentamicin resistance. Colonies were chosen for gentamicin sensitivity and loss of sucrose (5%) sensitivity, which typically indicates a double-crossover event and successful gene replacement, yielding the Δ*fliC* mutant. Deletion of *fliC* was confirmed by PCR. Primers are listed in Table S2.

A similar strategy was employed to construct the Δ*flgE* mutants. Briefly, PCRs were performed to amplify sequences upstream and downstream of *flgE*. The upstream fragment was amplified from PAO1 genomic DNA using primers D-*flgE*-up-F and D-*flgE*-up-R, while the downstream fragment was amplified with primers D-*flgE*-down-F and D-*flgE*-down-R. The vector pEX18Ap was assembled with the upstream and the downstream regions of the *flgE* gene using MultiF Seamless Assembly Mix, yielding pEX18Ap::*flgEUD*. A 1.8 kb gentamicin resistance cassette was cut from pPS858 with BamHI and then cloned into pEX18Ap::*flgEUD*, yielding pEX18Ap::*flgEUGD*. Primers are listed in Table S2.

### Construction of plasmids for gene complementation

For the construction of plasmid p-*fliC* (i.e., pAK1900::*fliC*), an ∼1.5 kb DNA fragment covering *fliC* was amplified from *P. aeruginosa* PAO1 genomic DNA with primers *fliC*-comp-F and *fliC*-comp-R by PCR. The vector pAK1900 was assembled with the PCR product of the *fliC* gene using MultiF Seamless Assembly Mix, yielding p-*fliC* (*38*). Primers are listed in Table S2.

A similar strategy was used to construct p-*flgE*. For the generation of plasmid p-*flgE* (i.e., pAK1900::*flgE*), an ∼1.4 kb DNA fragment containing *flgE* was amplified from *P. aeruginosa* PAO1 genomic DNA with primers *flgE*-comp-F and *flgE*-comp-R by PCR. The vector pAK1900 was assembled with the PCR product of the *flgE* gene using MultiF Seamless Assembly Mix, yielding p-*flgE*. All the constructs were verified by DNA sequencing. Primers are listed in Table S2.

### CHO cells and Plasmid transfection

The *Kcnq4* cDNAs were kindly provided by M. Shapiro (University of Texas Health Science Center, San Antonio, TX, USA). The *Tlr5* plasmid (pcDNA3.1-CMV-Tlr5-3×FlAG-hGHpolyA-EF1a-EGFP, pcDNA3.1-CMV-MCS-3×FlAG-hGHpolyA-EF1a-EGFP for vehicle control) were purchased from OBiO Technology (Shanghai, China). Chinese hamster ovary (CHO) cells were cultured in DMEM/F-12 medium (Gibco, Carlsbad, CA, USA) supplemented with 10% fetal bovine serum (FBS). For transient transfection, CHO cells were co-transfected with a total of 4 μg of cDNA (at a 1:1 ratio of *Kcnq4* to *Tlr5*) using the Lipofectamine 2000 reagent (Invitrogen, Carlsbad, CA, USA) according to the manufacturer’s instructions. Electrophysiological recordings were conducted 24-36 h after transfection.

### Method details

#### Bacterial strains and culture

For each experiment, the corresponding bacterial strains were streaked onto LB agar plates supplemented with carbenicillin (150 μg/mL) and grown overnight at 37°C. Single colonies were then picked and cultured overnight (approximately 14 h) in LB liquid medium containing carbenicillin (150 μg/mL) at 37°C with shaking at 210 rpm. Bacterium growth was monitored by measuring the optical density at 600 nm (OD600). Cultures at an OD600 of 5-6 (corresponding to 5-6 × 10⁸ CFU/mL) were used for subsequent mouse behavioral experiments.

### Bacterial growth curve measurement

The bacterial growth was measured in 96-well plates (Costar, Corning Incorporated, Corning, NY, USA). Overnight cultures of *P. aeruginosa* strains grown in LB medium were diluted into 10 mL of fresh LB in 50 mL tubes to an initial OD600 of approximately 0.05. Aliquots (100 μL) of the diluted PAO1 cultures were distributed into 96-well plates and overlaid with 70 μL of liquid paraffin to prevent evaporation. The plates were then incubated at 37°C, and OD600 was recorded at regular intervals by Synergy 2 Multi-Mode Microplate Reader (BioTek, Winooski, VT, USA).

### Swimming motility assays

Swimming motility of *P. aeruginosa* strains was performed as previously described (*39*). Swimming plates were prepared with LB medium solidified with 0.25% agar. Overnight cultures of *P. aeruginosa* strains grown in LB medium were washed three times and resuspended in PBS to an initial OD600 of approximately 0.1. Aliquots (3 μL) of the bacterial suspensions were spotted onto the center of swimming plates. The plates were allowed to dry at room temperature for 30 min and then incubated at 37°C for 20 h. Motility was evaluated by measuring the diameter of the bacterial growth zone.

### Transmission electron microscope (TEM) analysis

*P. aeruginosa* strains were cultured overnight in 5 mL of LB medium at 37°C with shaking. Bacterial samples were then imaged using a Talos L120C transmission electron microscope operating at 120 kV (Thermo Fisher Scientific, Waltham, MA, USA) as previously described (*40*).

### Mechanical itch (Alloknesis) test

The fur on the nape back of experimental mice was shaved before test. Mice were acclimated in a red recording chamber with a removable mesh cover for five consecutive days. On testing days, mice received mechanical stimulus for 1 s delivered using a von Frey filament (North Coast Medical, bending force: 0.07 g) at five randomly selected points, with each point 3 times. Stimulations at each point had an interval of 10 s. The scratching response of the hind paw toward the poking site was considered a positive response. Every three positive responses are counted as one point, with a maximum score of five. The scratching number at each point was summed into the final score for comparison.

### Spontaneous itch

After a three-day habituation period following dorsal shaving, each mouse was placed individually in a red recording chamber and video-recorded for 1.5 h. The first 30 min served as adaptation, scratching bouts during the subsequent 60 min were quantified as spontaneous itch. A scratching bout was defined as previously described (*12*).

### Acute itch test

Mice were habituated in the behavioral testing apparatus for 30 min. Then corresponding reagents including flagellin (Yeasen, Cat# 92531ES10; 0.3 or 0.9 μg/20 μL) or dechloroclozapine (DCZ; Sparkjade, Cat# SJ-MX3792; 1 μmol/20 μL) were intradermally injected into the nape of the back in mice. Immediately following injection, the mice were video-recorded for 30 min, and the number of scratching bouts during this period was quantified. Scratching was defined according to the same criteria used for spontaneous itch. Test the mechanical itch score at the injection site edge 30 min after recording acute itch.

### Mechanical pain

Mice were habituated on an elevated platform with a mesh floor for 30 min. The plantar side of the hindpaw was stimulated with calibrated von Frey filaments of defined forces (0.008-2 g). The 50% paw withdrawal thresholds were calculated using the Up-Down method to assess mechanical pain baseline (*10*). Then corresponding reagents including flagellin (0.3 or 0.9 μg/10 μL) or DCZ (1 μmol/10 μL) were administered via intraplantar injection into the hindpaw. Mechanical pain thresholds were reassessed 1 h post-injection.

### Open field test (OFT)

Mice were gently placed in the center of a white, non-transparent plastic open-field arena (50 cm × 50 cm × 50 cm) and allowed to explore freely for 10 min. A video camera positioned directly above the arena was used to track the trajectory of each animal. The total distance traveled and the total time spent in the arena were analyzed using TSE VideoMot2 software (TSE Systems, Bad Homburg, Germany). The arena was cleaned with 70% ethanol after every trial to remove olfactory sense from the box.

### Rotarod test

Mice were placed on the rotarod fatigue tester (IITC Life Science, Woodland Hills, CA, USA) and allowed to acclimatize for 30 s as previously described (40719051). The rotarod was then activated, and the rotation speed was gradually increased from 5 to 40 rpm over a 5 min period. The latency to the first fall from the rod was recorded. Measure each mouse three times and take the average, with an interval of 30 minutes between each measurement. For three consecutive days prior to the formal test, mice were trained on the rotarod three times daily, with each training session lasting 5 min (*41*).

### Pharmacological silence of TLR5^+^ and TRPV1^+^ sensory fibers

To selectively block TLR5⁺ fibers, 20 μL solution containing 0.2% QX-314 (TargetMol, Cat# 5369-03-9) and flagellin (0.3 μg) was administered via intradermal injection into the nape of the neck, as previously described. For selective silencing of TRPV1^+^ fibers, 20 μL solution containing 0.2% QX-314 and capsaicin (10 μg; Macklin, Cat# 404-86-4) was injected. 20 μL solution of 0.2% QX-314 alone was intradermally injected into the nape of the neck. QX-314, flagellin, and capsaicin were dissolved in isotonic saline. Acute itch behavior was recorded for 30 min immediately following injection, after which mechanical itch was tested.

### In situ hybridization (ISH) and immunohistochemistry

Mouse samples for RNAscope and immunohistochemistry were prepared as previously described. Briefly, DRGs were dissected from mice and fixed in 4% paraformaldehyde (PFA) overnight at 4°C. Then, the DRGs were cryopreserved in a 30% sucrose solution for 24 h and embedded in O.C.T compound to cut into 12 μm-thick sections. In situ hybridization (ISH) was performed using the RNAscope Multiplex Fluorescent Reagent Kit (ACDBio, Cat# 323100) according to the manufacturer’s instructions. The RNA probes were complementary to target mouse *Tlr5* (ACDBio, Cat# 451601-C1), *Trpv1* (ACDBio, Cat# 313331-C3), *Trkb* (ACDBio, Cat# 423611-C3), and *Kcnq4* (ACDBio, Cat# 472271-C1) mRNAs. Fluorescent immunohistochemistry (IHC) was performed as previously described (*14*, *42*). DRG sections were incubated overnight at 4°C with the following primary antibodies: NF200 (Sigma, Cat# N4142; 1:300), TH (EMD Millipore, Cat# MAB318; 1:500), or Calbindin D-28k (Swant, Cat# CB38; 1:1000). Following three washes with phosphate-buffered saline (PBS), sections were stained with the corresponding secondary antibodies for 2 h at room temperature. Secondary antibodies used are Cy^TM^3-conjugated AffiniPure Donkey anti-Mouse lgG (Jackson, Cat# 715-545-151; 1:500) and Alexa Fluor 488-conjugated AffiniPure Donkey anti-Rabbit lgG (Jackson, Cat# 711-165-152; 1:500). Sections were then washed three times in PBS and mounted with DAPI (Sigma, Cat# MBD0020). All sections were imaged using a Leica inverted confocal microscope (Model SP5, Wetzlar, Germany). Image analysis was performed using Leica Application Suite X (LAS X) and Adobe Photoshop.

### Retrograde labeling

Alexa Fluor™ 488-conjugated Cholera Toxin Subunit B (CTB; 2 µL of 1% [wt/vol]) was injected subcutaneously into the dorsal skin of *Tlr5^Cre^;Ai9* mice. Mouse DRG tissues (vertebral levels L4-L6) were dissected 7 days post-injection for histochemical analysis.

### Fluorescence-Activated Cell Sorting

Fluorescence-activated cell sorting (FACS) was conducted in dissociated DRG cells isolated from *Tlr5^Cre^;Ai9* mice to obtain Tlr5^+^ neurons. To isolate and stain mouse TLR5^+^ neurons, DRGs were dissected from *Tlr5^Cre^;Ai9* mice and trimmed on ice. The ganglia were carefully digested in a solution containing 2.21 mg/mL dispase II (Sigma, Cat# D4693) and 2 mg/mL collagenase A (Sigma, Cat# 10103586001) at 37°C for 30-40 min with gentle agitation. The cell suspension was filtered through a 40 μm cell strainer to remove debris and large fibers, collected in a flow cytometry tube, and stored on ice. TLR5^+^ neurons, identified by *tdTomato*-derived red fluorescence (excitation 554 nm / emission 581 nm), were sorted using a BD Aria III flow cytometer with a target purity of ≥95%. Sorted cells were collected directly into 96-well plates pre-filled with 200 μL TRIzol Reagent (Invitrogen, Carlsbad, CA, USA) per well, at 5,000 cells per well, for subsequent total RNA extraction.

### Quantitative Real-Time PCR

Total RNA from sorted TLR5^+^ neurons was extracted with TRIzol Reagent following the manufacturer’s instructions. The NanoDrop spectrophotometer (Thermo Fisher Scientific, Waltham, MA, USA) was employed for RNA quality control. Linear acrylamide (12 μL) was added to facilitate RNA precipitation. mRNA was reverse-transcribed to cDNA following the instruction of the Maxima H Minus First Strand cDNA Synthesis Kit (Thermo Fisher Scientific, Waltham, MA, USA). For quantitative real-time PCR (qPCR), gene-specific primers and cDNA were combined with Power SYBR™ Green PCR Master Mix (Bio-Rad, Hercules, CA, USA), and reactions were performed on a QuantStudio Real-Time PCR instrument (Applied Biosystems™ 7500; Thermo Fisher Scientific, Waltham, MA, USA). Primers are listed in Table S2.

### Whole-cell electrophysiology

Electrophysiological experiments were performed in cultured large-diameter DRG neurons or CHO cells using an EPC-10 patch-clamp amplifier and Patchmaster 2.2 software (HEKA Electronik, Lambrecht, Germany). Signals were filtered at 2 kHz and digitized at 10 kHz. Patch pipettes were pulled from borosilicate glass capillaries (World Precision Instruments, Sarasota, FL, USA) with an electrode resistance of 2-5 MΩ. Data analysis was performed using pClamp software (version 10.4; Molecular Devices, Sunnyvale, CA, USA).

M-type K^+^ currents in DRG neurons were recorded in voltage-clamp mode. The pipette (internal) solution contained the following (in mM): 140 KCl, 20 HEPES, 5 EGTA, 3 MgCl2, and 3 ATP-Na2 (pH adjusted to 7.3 with KOH). The bath (external) solution consisted of (in mM): 120 Choline-Cl, 3 KCl, 5 HEPES, 11 glucose, 2.5 CaCl2, and 1.2 MgCl2 (pH adjusted to 7.3 with NaOH). Neurons were held at -20 mV and followed by a voltage pulse to -60 mV. M-type K^+^ currents were measured as the deactivating tail currents at -60 mV.

KCNQ4 currents in CHO cells were recorded using the whole-cell voltage-clamp technique. The pipette solution contained (in mM): 140 KCl, 10 HEPES, 5 EGTA, and 1 MgCl2 (pH adjusted to 7.3 with KOH). The bath solution consisted of (in mM): 140 NaCl, 5 KCl, 10 HEPES, 10 glucose, 2 CaCl2, and 1 MgCl2 (pH adjusted to 7.3 with NaOH).

Current-clamp recording was performed to record the action potential firing properties of DRG neurons as previously described (*43*). Patch pipettes were filled with solution contained (in mM): 140 KCl, 0.5 EGTA, 5 HEPES, and 3 Mg-AT (pH adjusted to 7.3 with KOH). The bath solution contained (in mM): 140 NaCl, 3 KCl, 2 MgCl2, 2 CaCl2, and 10 HEPES (pH adjusted to 7.3 with NaOH). A whole-cell configuration was obtained in voltage-clamp mode, and the recording was performed after switching to current-clamp mode. Resting membrane potential (RMP) was assessed in current-clamp mode with zero current injection for at least 2 min. Current threshold was determined by the first action potential firing evoked by a series of depolarizing current injections that increased in 10 pA increments. Action potential (AP) half-width was measured at 50% of action potential amplitude.

### Quantification and statistical analysis

Results are expressed as mean ± SEM. Statistical analysis was performed in Prism 9 (GraphPad). Data from both male and female genders were pooled together for further analysis with unpaired two-tailed Student’s *t-*test for two groups. For experiments involving two groups across multiple time points, two-way repeated measures ANOVA followed by Šídák’s multiple comparisons test was used. For experiments comparing three or more groups at a single time point or across sessions, one-way or two-way ANOVA followed by Tukey’s post-hoc test was applied to determine differences between all possible pairs. A threshold of *P* < 0.05 was accepted as statistically different and *P* > 0.05 was considered non-significant. Statistical details including statistical tests used, number of samples (n), and P-values are reported in figure legends.

## Acknowledgements

The authors are grateful to Prof. Shenbin Liu (Fudan University Brain Science Research Institute) for generously sharing *Kit^W-sh/W-sh^* mice and to M. Shapiro (University of Texas Health Science Center at San Antonio) for providing the Kcnq4 cDNA constructs used in this work.

## Author contributions

Methodology: Z.P., H.D., H.L., Q.W., Y.W., and Q.Z. Investigation: Z.P., H.D., H.L., Q.W., Y.W., Q.Z., G.Z., X.H., Z.C., L.Z., and T.W. Formal analysis: Z.P., H.D., G.Z., X.H., Z.C., L.Z., and T.W. Visualization: Z.P., H.D. Funding acquisition: F.L., J.F., and H.Z. Supervision: F.L., J.F., H.Z., L.L., and Z.G. Writing—original draft: Z.P. and H.D. Writing—review & editing: F.L., J.F., H.Z., Y.C., Z.P., and H.D. All authors reviewed and approved the final manuscript.

## Data, code, and materials availability

All data and code needed to evaluate and reproduce the results in the paper are present in the paper and/or the Supplementary Materials. This paper does not report original code. All bacterial mutant strains generated in this study are available upon request. Other data used in this study reported in this paper is available upon request.

