## Supplemental Figs.1 to 6, Supplemental Tables 1 and 2 for "A microbial-sensory axis drives *Pseudomonas aeruginosa*-induced mechanical itch"

Science Advances

AAAS

Zhaohua Pang *et al.*

**This PDF file includes:**

Figs. S1 to S6
Tables S1 and S2

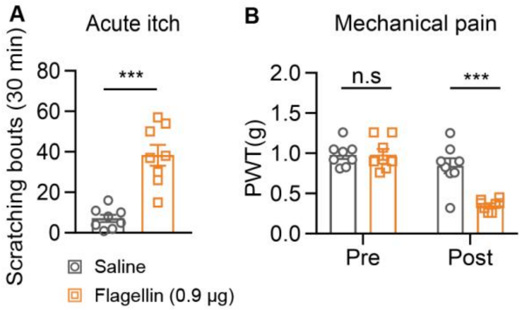


**Fig. S1. High-dose flagellin recruits nocifensive and canonical pruriceptive modalities. Related to Figure 2.** (A-B) Acute spontaneous itch bouts (A) and mechanical pain thresholds (B) evaluated following intraplantar injection of saline or 0.9 μg purified flagellin in wild-type mice (n = 8 mice per group). Data are presented as mean ± SEM. Statistical analyses: unpaired two-tailed Student's *t*-test (A); two-way repeated-measures ANOVA followed by Šídák's multiple comparisons test (B). n.s., not significant; ***$P < 0.001$.

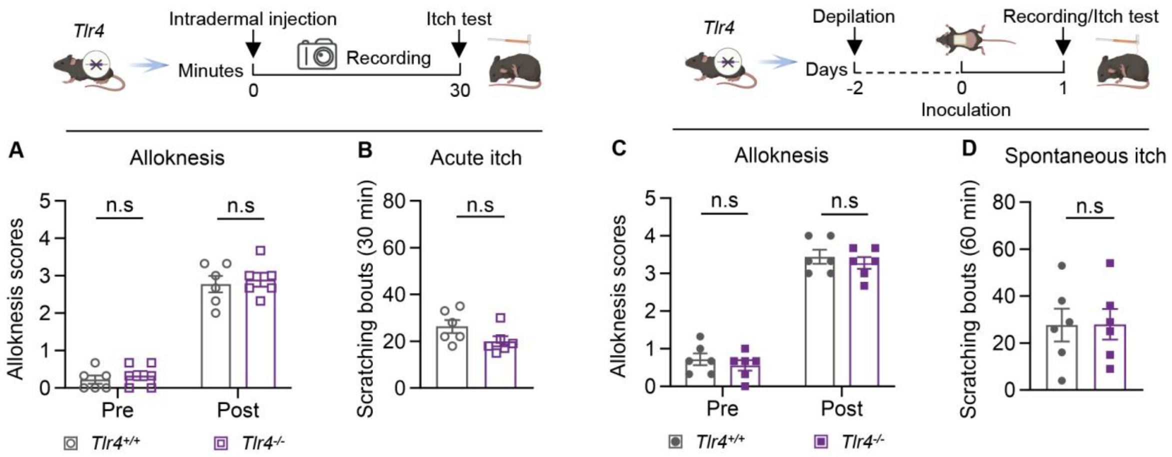


**Fig. S2. Global TLR4 deficiency does not attenuate flagellin- or PAO1-evoked alloknesis. Related to Figure 3.** (A-B) Mechanical alloknesis scores (A) and acute spontaneous itch bouts (B) following intradermal injection of 0.3 µg flagellin in *Tlr4*$^{-/-}$ (n = 8 mice) and *Tlr4*$^{+/+}$ littermates (n = 6 mice). (C-D) Mechanical alloknesis scores (C) and spontaneous itch bouts (D) evaluated before and after epicutaneous PAO1 exposure in *Tlr4*$^{-/-}$ and *Tlr4*$^{+/+}$ littermates (n = 6 mice per genotype). Data are presented as mean ± SEM. Statistical analyses: two-way repeated-measures ANOVA followed by Šídák's multiple comparisons test (A, C); unpaired two-tailed Student's *t*-test (B, D). n.s., not significant.

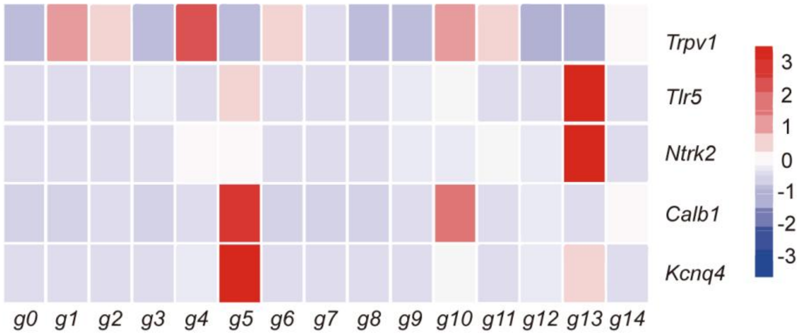


**Fig. S3. Transcriptomic mapping of *Tlr5* and *Kcnq4* across murine DRG neuronal subpopulations. Related to Figure 4.** Single-cell RNA-seq expression heatmap [re-analyzed from delineating the selective enrichment of *Tlr5*, *Calb1*, *Ntrk2*, *Trpv1*, and *Kcnq4* transcripts across discrete naïve mouse dorsal root ganglion (DRG) neuronal clusters (g0-g14)].[27] Color scale denotes row-normalized expression levels (blue, low; red, high).

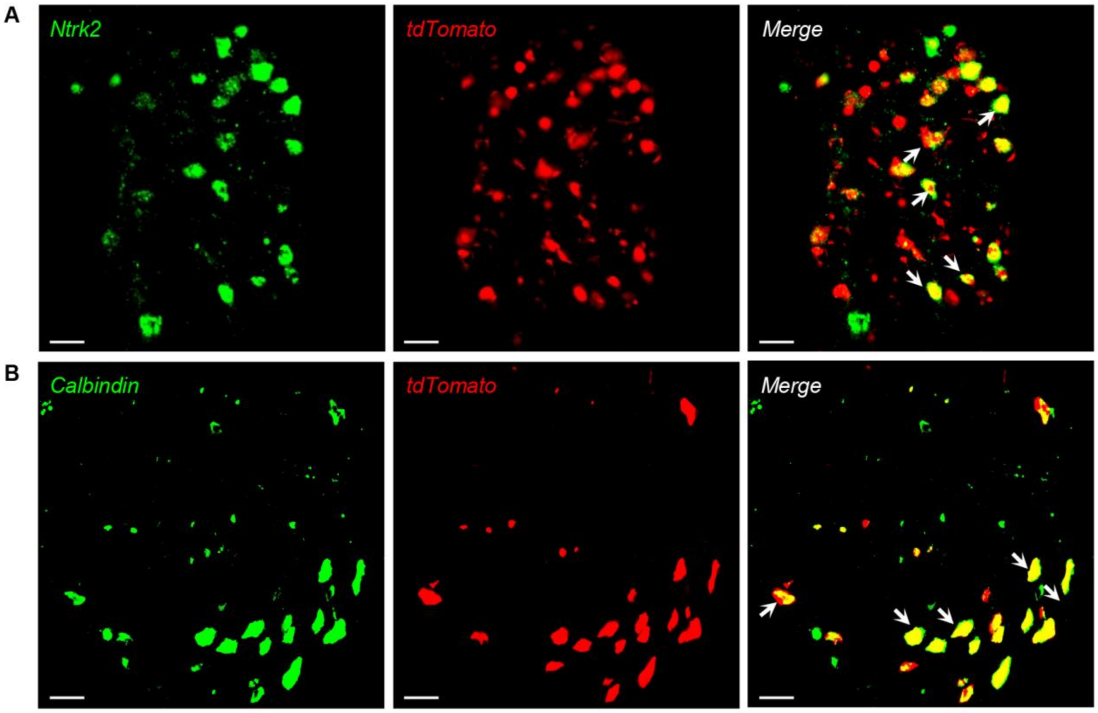


**Fig. S4. Validation of genetic reporter fidelity for LTMR subpopulations. Related to Figure 4.** (A-B) Representative whole-mount DRG co-labeling demonstrating the strict spatial congruence between (A) endogenous *Ntrk2* mRNA (green, ISH) and *Ntrk2*$^{CreER}$-driven tdTomato fluorescence (red), and (B) endogenous Calbindin protein (green, IF) and *Calb1*$^{dgCre}$-driven tdTomato fluorescence (red). White arrows indicate co-expressing neurons. Scale bar, 50 μm.

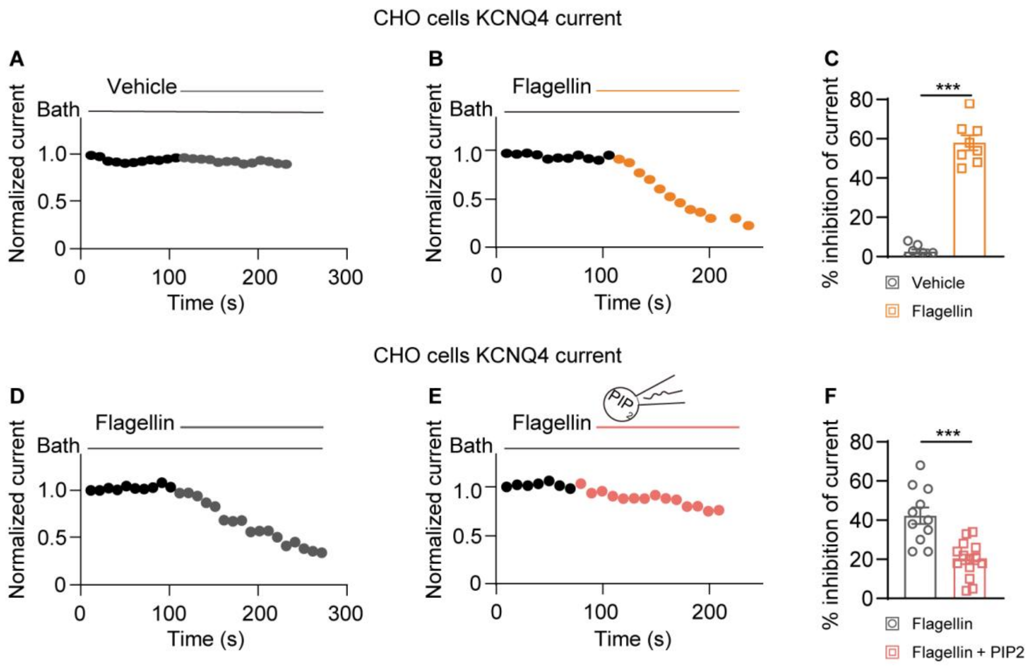


**Fig. S5. Intracellular PIP2 supplementation prevents flagellin-mediated M-current suppression *in vitro*. Related to Figure 6.** (A-C) Representative voltage-clamp traces of M-currents (A, B) and corresponding inhibition quantification (C) in KCNQ4-expressing CHO cells acutely exposed to vehicle or 50 nM flagellin. (D-F) Representative M-current traces (D, E) and inhibition quantification (F) demonstrating that intracellular dialysis of PIP2 (200 μM) abolishes flagellin-induced M-current suppression. Data are presented as mean ± SEM. Statistical analyses: unpaired two-tailed Student's *t*-test (C, F). $^{***}P < 0.001$.

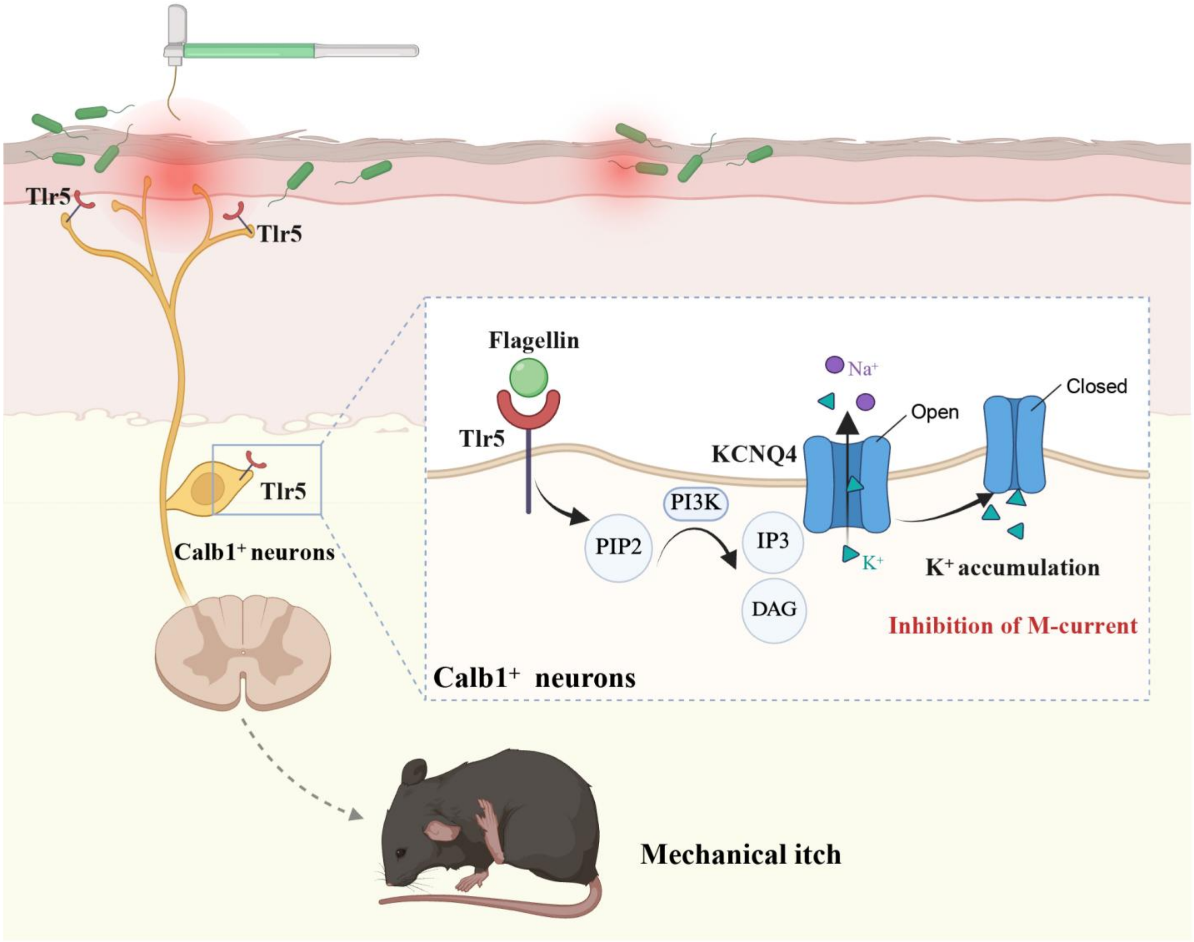


**Fig. S6. Proposed mechanistic model of the flagellin-TLR5-KCNQ4 axis driving infection-associated alloknesis.**

During cutaneous *P. aeruginosa* infection, structural bacterial flagellin directly engages TLR5 on the peripheral terminals of Calb1$^{+}$ Aβ rapidly adapting low-threshold mechanoreceptors (RA-LTMRs). This pathogen-receptor interaction triggers the intracellular depletion of membrane PIP2, which subsequently suppresses the activity of KCNQ4 voltage-gated potassium channels. The loss of KCNQ4-mediated M-currents dismantles the biophysical brake on these mechanosensors, intrinsically lowering their tactile activation threshold. Consequently, normally innocuous mechanical stimuli are aberrantly encoded and transmitted to downstream spinal itch circuits, culminating in pathological touch-evoked pruritus independent of canonical C-fiber nociceptive pathways.

**Table S1.**

Plasmids and bacterial strains and used in this study.

| **Plasmids or strains** | **Relevant characteristics** | **Source** |
|---|---|---|
| **Plasmids** | | |
| pEX18Ap | Gene replacement vector, mob$^{+}$*sacB*, Ap$^{r}$ | Jansons et al (*37*). |
| pPS858 | pBR322 derivative carrying a FRT-Gm cassette, Ap$^{r}$ | Jansons et al (*37*). |
| pAK1900 | *E. Coli-P. aeruginosa* shuttle cloning vector, Ap$^{r}$ Cb$^{r}$ | Hoang et al (*38*). |
| p-*fliC* | PAK1900 derivative carrying *fliC* (PA1092) on a *c*. 0.8 kb *HindIII/BamHI* fragment in same orientation as *plac* | This study |
| p-*flgE* | PAK1900 derivative carrying *fliE* (PA1080) on a *c*. 0.8 kb *HindIII/BamHI* fragment in same orientation as *plac* | This study |
| pEX18Ap::*fliCUGD* | pEX18Ap derivative, for replacing *fliC* gene with a gentamicin resistance cassette from plasmid pPS858 | This study |
| pEX18Ap::*flgEUGD* | pEX18Ap derivative, for replacing *flgE* gene with a gentamicin resistance cassette from plasmid pPS858 | This study |
| ***P. aeruginosa* strains** | | |
| PAO1 | Wild type, WT | Stover et al (*36*). |
| Δ*fliC* | PAO1 derivative with a gentamicin resistance cassette replaced the *fliC* gene | This study |
| Δ*flgE* | PAO1 derivative with a gentamicin resistance cassette replaced the *flgE* gene | This study |
| ***E. coli* strains** | | |
| DH5α | *endA hsdR17 supE44 thi-1 recA1 gyrA relA1*Δ (*lacZYA-argF*) *U169 deoR* (*φ80dlac*Δ(*lacZ*)*M15*) | This laboratory |
| S17 λ-pir | *recA thi pro hsdR*$^{-}$*M*$^{+}$ RP4-2-Tc::Mu Km::Tn7 λpir (Tp$^{r}$ St$^{r}$) | This laboratory |

Ap$^{r}$, ampicillin resistance; Cb$^{r}$, carbenicillin resistance; Gm$^{r}$, gentamycin resistance; Tp$^{r}$, trimethoprim resistance; St$^{r}$, Chromosomal streptomycin resistance.

**Table S2.**

Primer sequences used in this study.

| Organism | Amplicon | Primer (5'→3') | Sequence |
|---|---|---|---|
| **Bacteria** | D-*fliC*-up | Forward | ACAGCTATGACCATGATTACTTTCGTGGTCTCGACCGAAC |
| | | Reverse | AGGCTCAGGATCTAGAGTGTTGACTGTAAGGGCCAT |
| | D-*fliC*-down | Forward | CAGTCAACACTCTAGATCCTGAGCCTGCTGCGCTAA |
| | | Reverse | GTAAAACGACGGCCAGTGCCTTGAACTTGGCATTGGCCGG |
| | D-*flgE*-up | Forward | ACAGCTATGACCATGATTACAGGACCAGAGTTCGCTGAT |
| | | Reverse | TTGATGATGGTCTAGAAGGCCGATGTTGAAACTCAT |
| | D-*flgE*-down | Forward | ACATCGGCCTTCTAGACCATCATCAACCTGCGCTGA |
| | | Reverse | GTAAAACGACGGCCAGTGCCATTGACGGGACAGCGAGAGG |
| | *fliC*-comp | Forward | GACACTATAGAATACTCAGGTCCTTTGGAGGAAATCACC |
| | | Reverse | AATTCGAGCTCGGTACCCGTACCGCGTGAGTGACCGTT |
| | *flgE*-comp | Forward | GACACTATAGAATACTCGTTTCCGGCAAGGAGCTATC |
| | | Reverse | AATTCGAGCTCGGTACCCGGTCATCAGCGCAGGTTGATG |
| **Mouse** | *β*-actin | Forward | GTGACGTTGACATCCGTAAAGA |
| | | Reverse | GCCGGACTCATCGTACTCC |
| | *Kcnq2* | Forward | GGGGCCCAACAATAACG |
| | | Reverse | TTTCTCCACCTTCCCA |
| | *Kcnq3* | Forward | CGCGCTTGTGTTCCTGATTG |
| | | Reverse | CAGCCCAGATCCTCAAAGCA |
| | *Kcnq4* | Forward | TATGGTGACAAGACGCCACAT |
| | | Reverse | GCTTCTCAAAGTGCTTCTGCC |
| | *Kcnq5* | Forward | ATTGGCTATGGAGACAAAACACC |
| | | Reverse | CGGTGCTGCTCCTGTACTTTT |
